# Metabolic and functional impairment of CD8^+^ T cells from the lungs of influenza-infected obese mice

**DOI:** 10.1101/2020.04.19.047282

**Authors:** William D. Green, Abrar E. Al-Shaer, Qing Shi, Nancie J MacIver, Melinda A. Beck, Saame Raza Shaikh

**Author notes:** These authors contributed equally to this work (William D. Green, Abrar E. Al-Shaer). Correspondence: Saame Raza Shaikh, 2203 McGavran Greenberg, University of North Carolina at Chapel Hill, Chapel Hill NC 27599.

## Abstract

**Background:** Obesity increases influenza disease risk in millions of adults worldwide. In this study, we investigated the effect of diet-induced obesity on pulmonary CD8^+^ T cell metabolism and function as a mechanism of impairment.

**Methods:** Male C57BL/6J mice were fed either control (10% kcal/g) or high-fat (60% kcal/g) diet. Sub-lethal A/PR/8/34 influenza virus infection generated a robust pulmonary immune response. T cell metabolism and function were assessed at day 10 and day 24 post infection.

**Results:** At day 10 post infection, CD8^+^ T cells from obese mice had impaired oxidative and glycolytic metabolism, greater fatty acid uptake, and decreased effector populations and cytokine production. At infection resolution, obese mice had lower numbers of naïve and central memory CD8^+^ T cell populations in the lungs.

**Conclusion:** Diet-induced obesity increases influenza virus pathogenesis through CD8^+^ T cell mediated metabolic reprogramming resulting in suppressed effector CD8^+^ T cell function.

**Summary:** Diet-induced obesity impairs the metabolism of pulmonary CD8^+^ T cells resulting in reduced effector CD8^+^ T cells and cytokine production following primary influenza infection.

## INTRODUCTION

Influenza virus causes significant global morbidity and mortality each year. Traditionally, high-risk groups for increased mortality from influenza include children under the age of five, the elderly, pregnant women and individuals with compromised lung or heart function. Importantly, obesity is also recognized as an independent risk factor for influenza infection, resulting in increased rates of hospitalization, complications such as pneumonia, and increased mortality (1–4). This is significant, because as of 2014, the global burden of obesity outnumbered underweight for the first time in human history, signifying an epidemic shift in the burden of increased weight on human health (5). This intersection between a chronic disease condition (obesity) and an acute infectious disease (influenza) represents a confluence of public health problems for the 21^st^ century. Currently, 10% of the global population, including 37% of the U.S. adult population, are obese, thereby putting them at increased risk of influenza associated morbidity and mortality.

Immune protection to influenza infection and vaccination relies on a competent innate and adaptive immune cell response, both of which are impaired in the obese host (6–15). These impairments are thought to be responsible for the observed increased risk to both seasonal and pandemic influenza in obese adults (2, 16). In humans, obese adults have greater rates of hospitalization and severe influenza (3, 17) and a sustained duration of influenza viral shedding (18). Our own studies have found influenza vaccination of obese adults, when compared with lean adults, results in a suboptimal vaccine response. When stimulated with influenza antigens, influenza specific CD4^+^ and CD8^+^ T cells from vaccinated obese adults have reduced expression of activation markers and impaired function (10). Alarmingly, despite having generated a “protective” level of flu-specific antibody, impaired T cell function is associated with vaccinated obese adults having twice the risk of influenza infection or an influenza-like illness (19).

Animal models of obesity mirror findings in humans. Diet-induced obese (DIO) mice demonstrate increased mortality to primary (14) and secondary (6) influenza infections, decreased influenza specific memory T cell populations (7), and suppressed memory T cell function (6). However, the underlying mechanism(s) by which obesity drives T cell dysfunction remains unclear.

Obesity is primarily a metabolic disease, and T cells rely on specific metabolic programs for their function. Typically, naïve and memory T cells utilize oxidative metabolism of fatty acids and glucose-derived pyruvate to drive and support immune surveillance and survival (20, 21). Activated effector T cells display predominately glycolytic and glutamine oxidative metabolism to fuel daughter cell generation and effector function in response to antigen (22–24). In a model of secondary influenza infection, in which obese mice had functionally impaired memory CD4^+^ and CD8^+^ T cells, splenic T cells were metabolically impaired both prior to infection and at 7 days post secondary infection (25). However, the spleen is not the most appropriate site to test whether impairments in T cell metabolism affect T cell function, as the lungs are the primary location of influenza infection. Therefore, the objective of this study was to determine if obesity promotes pulmonary T cell metabolic reprogramming during a primary influenza infection.

Using a DIO model, we infected lean and obese mice with a sublethal dose of mouse adapted influenza virus (A/PR/8/34). Here, for the first time, we report obesity impairs the metabolic reprogramming of effector CD8^+^ T cells within the lungs during influenza infection. Our results show CD8^+^ T cells from obese mice have suppressed glycolytic and oxidative metabolism with increased uptake of neutral fatty acid, concomitant with reduced CD8^+^ T cell effector populations and reduced production of the inflammatory cytokine, interferon gamma (IFN-γ). These results provide the first *in vivo* evidence of metabolic reprogramming in obesity during influenza viral infection in the lungs, suggesting obesity-induced impairment of CD8^+^ T cell metabolism influences influenza virus pathogenesis.

## MATERIALS AND METHODS

### Mice and Diets

C57BL/6J Control (Stock No. 380056, “lean”) and C57BL/6J DIO (Stock No. 380050, “obese”) eighteen week old, male mice were obtained from The Jackson Laboratory (Bar Harbor, Maine, USA) and allowed one week of acclimation. Mice were group housed (4 per cage), maintained at ambient temperature, and given ad libitum access to food and water. Thirty-six lean mice were placed on a purified low fat control diet (D12450B, Research Diets, New Brunswick, NJ, USA) and thirty-six DIO mice were maintained on 60% kcal high fat diet (HFD) (Research Diets, D12492). All procedures were performed in accordance with protocols approved by the Institutional Animal Care and Use Committee (IACUC) at the University of North Carolina at Chapel Hill.

### Influenza Infection

Influenza infection followed previously described protocols (25). Briefly, mice were lightly anesthetized with isoflurane and infected intranasally with 50 μL of sterile PBS containing 0.0025 hemagglutination units (HAU) of A/PR/8/34 (ATCC® VR-95™, Manassas, VA, USA), a H1N1 influenza virus. Mice were weighed daily and sacrificed at day 0, 10 and 24-post infection.

### Flow Cytometry

Lungs were prepared for flow cytometry analysis as previously described (9, 26). The following antibodies were used: Alexa Fluor 700 Rat anti-mouse CD3 (17A2, BioLegend, San Diego, CA, USA), Pacific Blue Rat anti-mouse CD4 (GK1.5, BioLegend), Alexa Flour 594 Rat anti-mouse CD8a (53-6.7, BioLegend), FITC Rat anti-mouse CD62L (MEL-14, BioLegend), APC Rat anti-mouse Granzyme B (NGZB, eBioscience, San Diego, CA, USA), PE Rat anti-mouse Interferon gamma (XMG1.2, eBioscience), PE Rat anti-mouse CD62L (MEL-14, BioLegend), APC Rat anti-mouse CD44 (IM7, BioLegend), APC-Cy7 Armenian Hamster CD69 (H1.2F3, BioLegend), APC Rat anti-mouse CD11a (M17/4, BioLegend), PE Armenian Hamster anti-mouse CD103 (2E7, BioLegend), Zombie Aqua Fixable Viability dye (BioLegened), UltraComp eBeads compensation beads (ThermoFisher, Waltham, MA, USA), purified Rat anti-mouse CD16/CD62 FcBlock (2.4G2, BD Bioscience). To determine fatty acid uptake, lung isolated single cells were incubated with 5μM BODIPY FL C_16_ (ThermoFisher) for 30mins at 37°C prior to extracellular staining. All samples were acquired on an Attune NxT flow cytometer (ThermoFisher), and data were analyzed using FlowJo v10 (Treestar).

### Extracellular Metabolic Flux Analysis

CD8^+^ T cells were isolated from the single cell suspension of mouse lungs at day 10 post PR8 infection using magnetic bead negative selection (Stemcell, Vancoover, Canada) in EasySep buffer (PBS + 2% FBS + 1mM EDTA). Isolated cells were counted using the Bio-Rad TC20 with trypan blue exclusion for viability. XFe96 cell culture microplates were treated with Cell-Tak™ (Corning, Corning, NY, USA) in 0.1M sodium bicarbonate to allow for cell adherence. CD8^+^ T cells were plated in non-buffered RPMI-1640 with freshly added 10mM glucose, 2mM glutamine, and 1mM pyruvate at 300,000 cells per well. Extracellular acidification (ECAR) and oxygen consumption rates (OCR) were determined using the Seahorse XFe96 Flux analyzer (Agilent, Santa Clara, CA, USA) at 37°C as previously described (27). OCR and ECAR were normalized to cell number.

### Statistical Analysis

Seahorse data were analyzed using custom-built R v3.4.4 and Python v2.7 scripts, linked here (https://github.com/abrar-alshaer/seahorse_analysis). We filtered outliers outside of the Q1/Q3±1.5*IQR range for technical or biological replicates. Parameters of the Mitochondrial Stress Test were calculated in our Python program as detailed in the Agilent Seahorse user guide. For statistical analyses, normality was tested using the Shapiro-Wilks test. If the data satisfied the assumption of normality, we utilized a Student T-test; otherwise we utilized the Wilcoxon rank-sum test. Details describing the statistical analysis and sample size of each experiment can be found in the figure legends. All statistical analysis was performed using GraphPad Prism 7 for Mac OS X, version 7.0c (GraphPad Software, Inc., La Jolla, CA) or R v3.4.4. All data was determined as significant by p<0.05.

## RESULTS

### Primary influenza infection elicits a robust pulmonary immune response

Following one week of acclimation, male C57BL/6J lean (n=36) and obese (n=36) mice were infected with a sublethal dose of A/PR/8/36 influenza virus (Figure 1A). Both lean and obese mice lost ~20% of their body weight by day 9 and 10 post-infection, with no difference in weight loss between lean and obese mice (Figure 1A, 1B). Lung and bronchoalveolar (BAL) fluid cell numbers in uninfected (Day 0), during infection (Day 10), and following immune resolution (Day 24) did not vary between lean and obese mice. As expected, both lean and obese mice had a dramatic increase in infiltrating immune cell numbers at day 10 (Figure 1C) relative to baselines (6.54 fold increase in lean; 7.91 fold increase in obese), demonstrating a robust primary immune response to influenza infection. Infection resolution resulted in significant decreases in cell numbers from day 10 peaks (51% decrease in lean, 49% decrease in obese), albeit with significantly higher cell numbers than baseline (2.65 fold greater in lean; 3.45 fold greater in obese).

**Figure 1.**
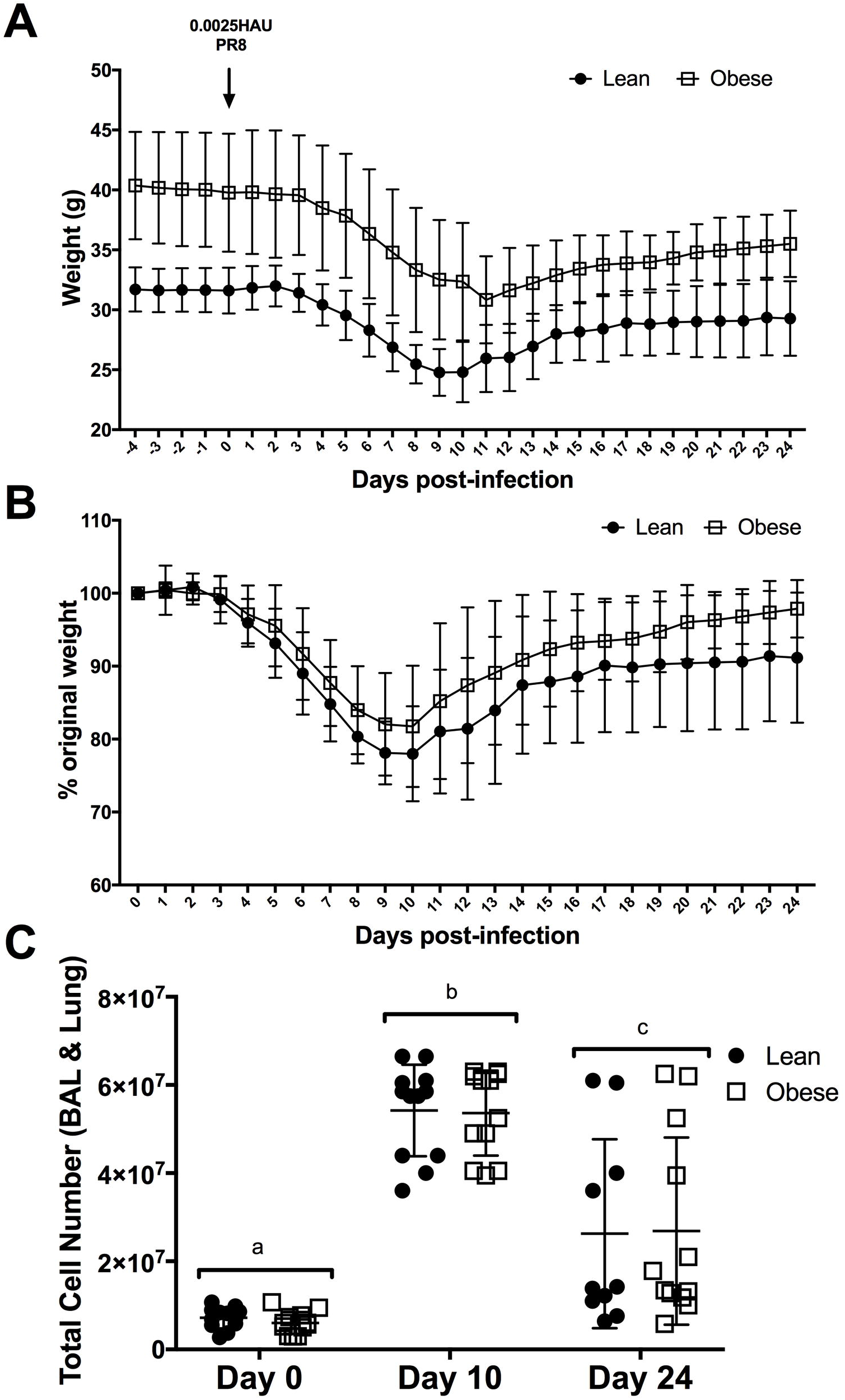
Sublethal influenza infection induces primary immune response in lungs of lean and obese mice. (A) Male, 18 week-old C57BL/6J mice were fed control (Lean, n=36) or high fat diet (Obese, n=36). After one-week acclimation, mice were weighed for four days before infection with 0.0025HAU PR8 influenza virus. Mice were maintained on diet for 24 days after infection. Body weights for measured daily. (B) Percent weight loss was calculated relative to original body weight at time of infection. (C) Lungs were harvested, digested, and homogenized in single cell suspension at each time point. Total cell number of combined bronchoaveolar lavage fluid and digested lungs from lean and obese mice at day 0, day 10, and day 24 post-influenza infection. Data represent mean ± SD (A,B) or each dot represents data obtained from one mouse with mean ± SD. Two-way ANOVA with Sidak’s multiple comparison was used to compare groups. Letter “a”, “b”, “c”, denote significance between time points.

### Obesity impairs CD8^+^ T cell metabolism in the lungs of influenza-infected mice

CD8^+^ T cells provide critical defense against influenza infection in the lungs. To establish if obesity alters the metabolic profile of these cells in the lungs during influenza infection, we isolated pulmonary CD8^+^ T cells from lean and obese mice at day 10-post infection. Oxidative and glycolytic metabolism was determined using an Agilent Seahorse extracellular flux analyzer by measuring oxygen consumption rate (OCR; a proxy for mitochondrial metabolism), and extracellular acidification rate (ECAR; a proxy for lactate production and glycolysis). Real time measurements of OCR (Figure 2A) and ECAR (Figure 2B) for pulmonary CD8^+^ T cells from influenza infected lean and obese mice show distinct metabolic differences day 10-post infection. Specifically, compared to pulmonary CD8^+^ T cells from influenza infected lean mice, pulmonary CD8^+^ T cells from obese mice had 30% lower basal OCR (Figure 2C) and 24.9% lower basal ECAR (Figure 2D).

**Figure 2.**
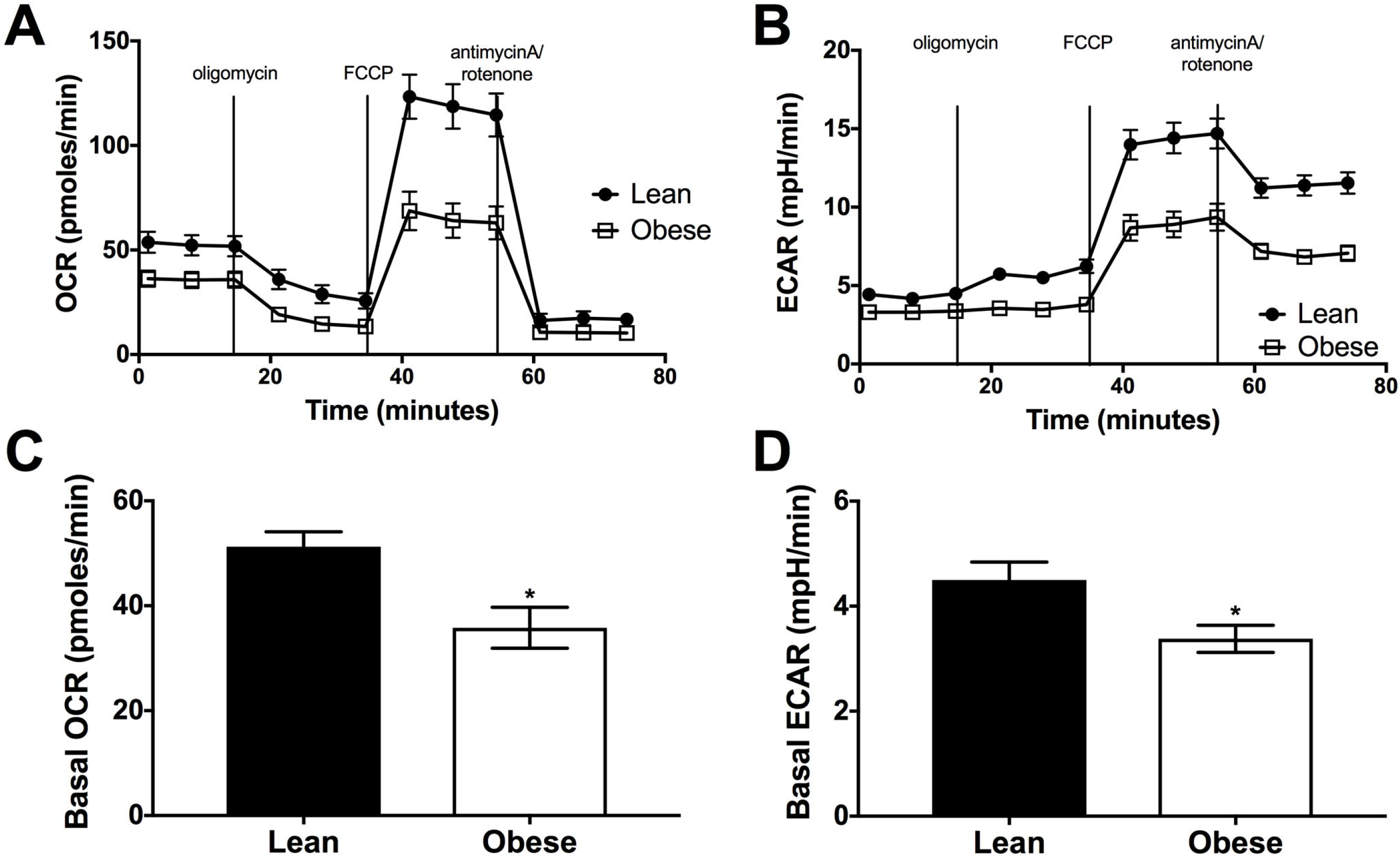
Obesity impairs CD8^+^ T cell metabolism in the lungs of influenza-infected mice. Lungs were harvested, digested, and homogenized into single cell suspension. Total CD8^+^ T cells from day 10-post influenza infected mice were isolated using negative selection magnetic bead separation. Cells were plated at 300,000 cells per well, and extracellular flux analysis was performed using the Agilent Seahorse XF_e_96 flux analyzer. (A) Oxygen consumption rates (OCR), a surrogate of glycolysis, and (B) Extracellular Acidification Rates (ECAR), a surrogate of mitochondrial respiration, were measured in response to the Mitochondrial Stress Test for CD8^+^ T cells from Lean (n=11) and Obese (n=12) mice. (C) Basal OCR and (D) Basal ECAR were determined as the last point before oligomycin injection. Data represents mean ± SEM and were normalized to cell number. Wilcoxon rank sum test was used to compare groups. *p<0.05, **p<0.01.

Maximal respiration, measured as the highest OCR following administration of the mitochondrial uncoupling agent carbonyl cyanide 4-(trifluoromethoxy) phenylhydrazone (FCCP), had 47.4% lower maximum mitochondrial respiration in pulmonary CD8^+^ T cells from obese mice compared to lean (Figure 3A). Similarly, compared to lean mice, CD8^+^ T cells from obese mice had 24.5% lower ATP production (Figure 3B) and 61.6% lower spare respiratory capacity (SRC; Figure 3C), a measure of the potential mitochondrial metabolic response to energy demand. Strikingly, the comparison of basal OCR versus basal ECAR before and after injection of oligomycin, an ATP-synthase inhibitor, showed impaired glycolytic potential of pulmonary CD8^+^ T cells from obese mice compared to lean (Figure 3D), signifying an inability of CD8^+^ T cells from influenza infected obese mice to use glycolysis over mitochondrial respiration.

**Figure 3.**
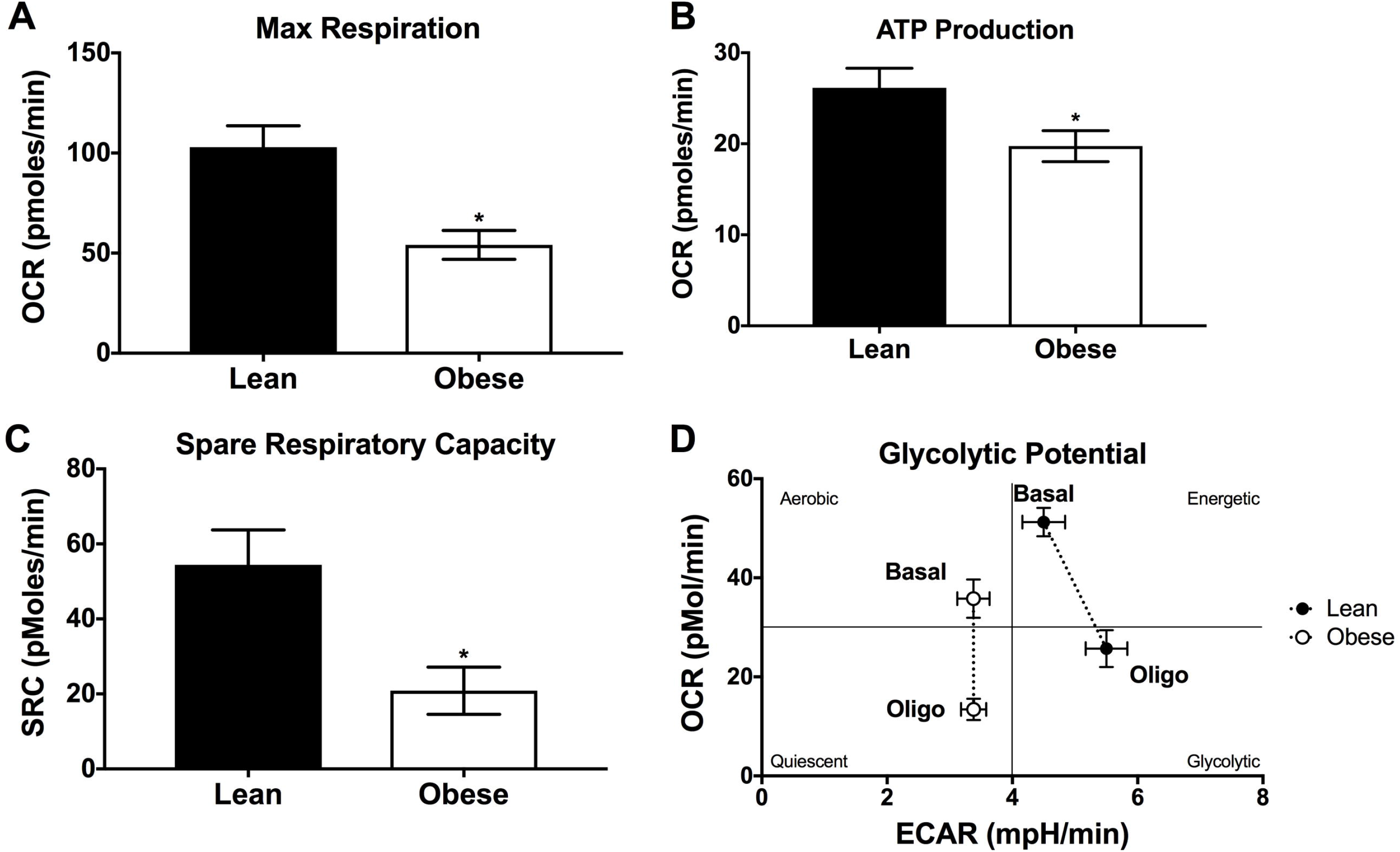
Pulmonary CD8^+^ T cell metabolic potential is impaired in influenza infected obese mice. Lungs were harvested, digested, and homogenized into single cell suspension. Total CD8^+^ T cells from day 10-post influenza infected mice were isolated using negative selection magnetic bead separation. Cells were plated at 300,000 cells per well, and extracellular flux analysis was performed using the Agilent Seahorse XF_e_96 flux analyzer and Mitochondrial Stress test for lean (n=11) and obese (n=12) mice. (A) Maximal respiration was measured as peak OCR following injection of 1.5μM FCCP. (B) Spare Respiratory Capacity was calculated as the difference in Max OCR minus Basal OCR. (C) ATP production was calculated as the difference between Basal OCR and the minimum OCR following injection of 0.5 μM oligomycin. (D) Glycolytic potential (OCR vs. ECAR) before and after injection of oligomycin. Data represents mean ± SEM and were normalized to cell number. Wilcoxon rank sum test was used to compare groups. *p<0.05.

### Greater fatty acid uptake in pulmonary CD8^+^CD44^−^CD62L^−^ T cells of influenza infected obese mice

Nutrient uptake supports metabolic reprogramming upon immune cell activation during infection. Previous foundational work has elegantly demonstrated the requirement of effector CD8+ T cells to use glycolytic metabolism to support effector cell production of cytokines, namely interferon gamma (IFN-γ) (28). Because pulmonary CD8^+^ T cells from obese mice demonstrated impaired glycolytic metabolism, we assessed if there was a change in fatty acid uptake in pulmonary CD8^+^ T cells between lean and obese mice. Obese mice had 35% reduction in frequency (Figure 4A) but not number (Figure 4B) of pulmonary effector CD8^+^ T cells (CD8^+^ CD44^−^ CD62L^−^) at day 10-post influenza infection. Representative (Figure 4C) and quantitative median fluorescent intensity (MFI; Figure 4D) of BODIPY FLC16 by effector CD8^+^ T cells within the lungs at day 10-post influenza infection demonstrates, effector CD8^+^ T cells from obese mice had 42.7% greater fatty acid uptake. No differences in fatty acid uptake were observed in pulmonary naïve, central memory, or effector memory CD8^+^ T cells at day 10-post infection (Figure S1A) or naïve, CM, EM, or residential memory at day 24 (Figure S1B). These results indicate obesity impairs activated effector T cell fuel uptake within the lungs during influenza infection.

**Figure 4.**
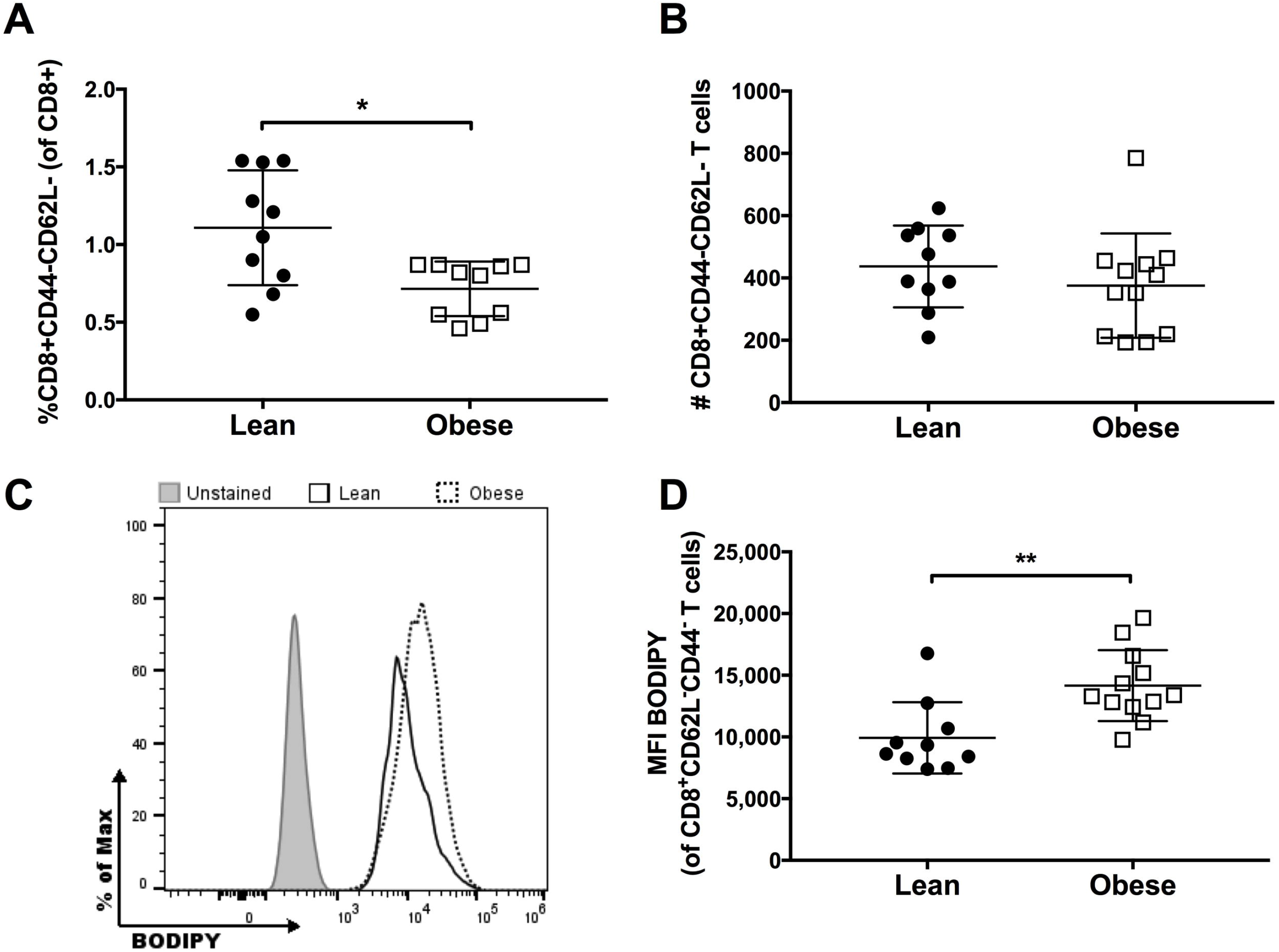
Obesity increases fatty acid uptake of effector CD8^+^ T cells in the lungs. Day 10 post-infection lungs were harvested, digested, and homogenized into single cell suspension and 1×10^6^ cells were stained for BODIPY FLC16 uptake by flow cytometry. (A) Frequency and (B) number of CD8^+^CD44^−^CD62L^−^ T cells from day 10 post-infection mice. (C) Representative histogram of BODIPY uptake normalized to mode by effector CD8^+^CD44^−^CD62L^−^ T cells from day 10 post-infection lean and obese mice. (D) Median fluorescence intensity (MFI) of effector CD8^+^CD44^−^CD62L^−^ T cells from lean (n=11) and obese (n=12) C57BL/6J male mice. Each dot represents data obtained from one mouse with mean ± SD. Wilcoxon rank sum test was used to compare groups. *p<0.05, **p<0.01.

### Obesity reduces pulmonary CD8^+^ T effector cells and cytokine production in influenza-infected mice

As both metabolism and nutrient uptake were altered in pulmonary CD8^+^ T cells, and since T cell metabolism drives T cell function (24, 29), we examined whether obesity also affected effector CD8^+^ T cells and cytokine production during influenza infection (Figure 5A). Similar to previous findings (6, 9, 25, 30), compared to lean mice, obese mice had significantly lower percent (Figure 5B) and number (Figure 5C) of granyzme B (GrB) positive and IFN-γ positive CD8^+^CD62L^−^ T cells at day 10 post-infection. This ~39% reduction in pulmonary effector CD8^+^ T cells occurred concomitant with reduced CD8^+^ T cell glycolytic capacity (Figure 3) and greater fatty acid uptake (Figure 4). Following influenza clearance at day 24-post infection, there was no difference in percent pulmonary effector CD8^+^CD62L^−^GrB^+^IFNg^+^ T cells (Figure 5B), albeit a non-significant reduction in number of CD8^+^CD62L^−^GrB^+^IFNg^+^ T cells (Figure 5C). Interestingly, at day 10, CD8^+^CD62L^−^GrB^+^IFNg^+^ T cells from obese mice also made significantly less IFN-γ (Figure 5D).

**Figure 5.**
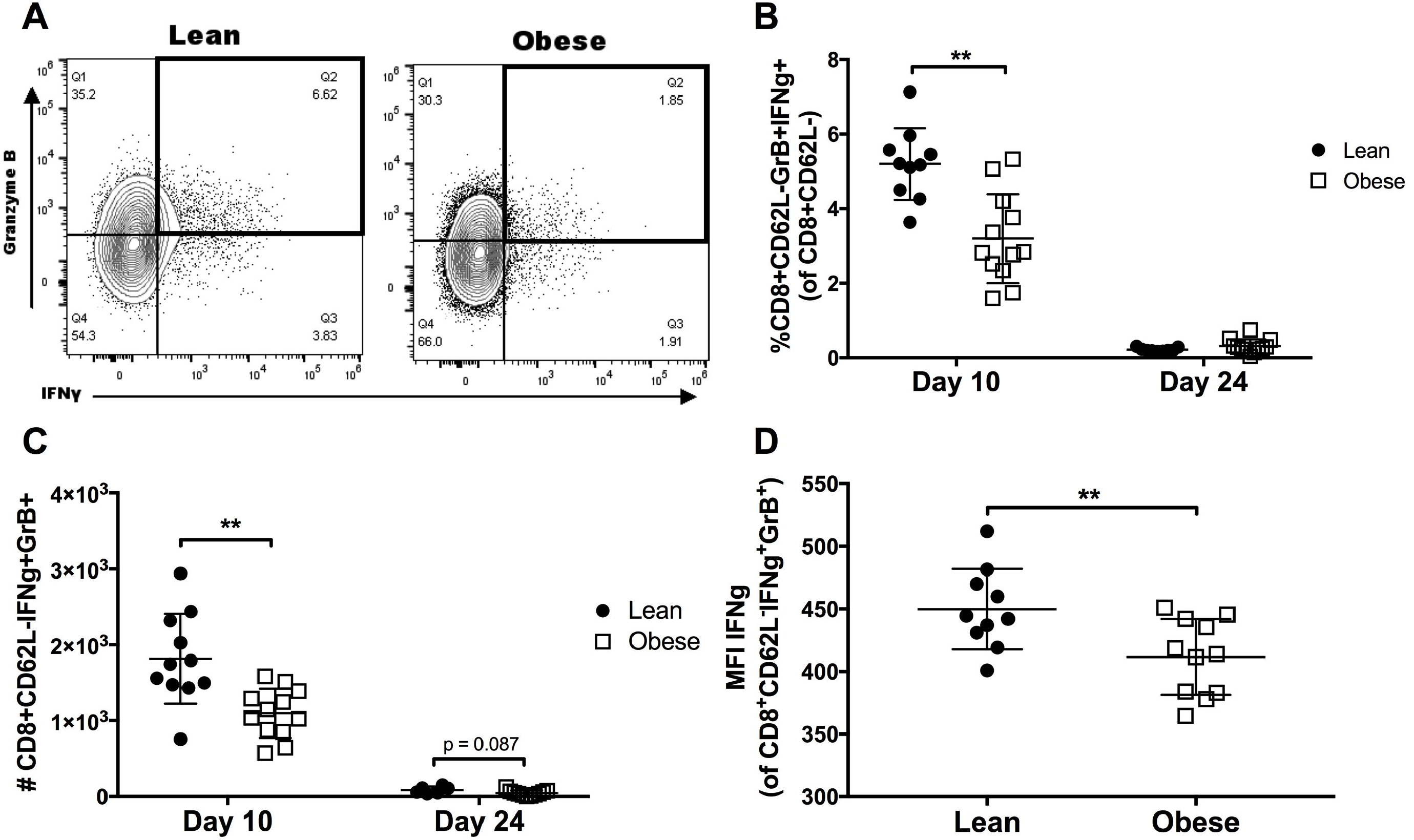
Obesity impairs CD8^+^ T cell production of effector cytokines in lungs of influenza-infected mice. Lungs were harvested, digested, and homogenized into single cell suspension and 1×10^6^ cells were stained for intracellular cytokines in effector T cells by flow cytometry. (A) Representative scatter plot of interferon gamma (IFN-γ) and granzyme B (GrB) double positive CD8^+^CD62L^−^ T cells at day 10-post infection from lean and obese mice. (B) Frequency and (C) cell number of CD8^+^CD62L^−^GrB^+^IFNγ^+^ T cells from day 10 and day 24 post influenza infection in lean (n=11) and obese (n=12) mice. (D) Median fluorescent intensity (MFI) of IFN-γ from CD8^+^CD62L^−^GrB^+^IFN ^+^ T cells at day 10 post influenza infection. Each dot represents data obtained from one mouse with mean ± SD. Wilcoxon rank sum test was used to compare groups. *p<0.05, **p<0.01.

Pulmonary effector CD8^+^ T cells making IFN-γ, but not GrB (Figure 6A), also had ~18% reduction in frequency (Figure 6B) and 20% reduction in number (Figure 6C) at day 10-post infection in obese mice. Again, no difference was observed at day 24-post infection, with a significant reduction in IFN-γ MFI at day 10-post infection (Figure 6D). Together, these data suggest impaired CD8^+^ effector T cell metabolism reduces the presence of effector CD8^+^ T cells as well as production of pro-inflammatory cytokines within the lungs of influenza infected obese mice.

**Figure 6.**
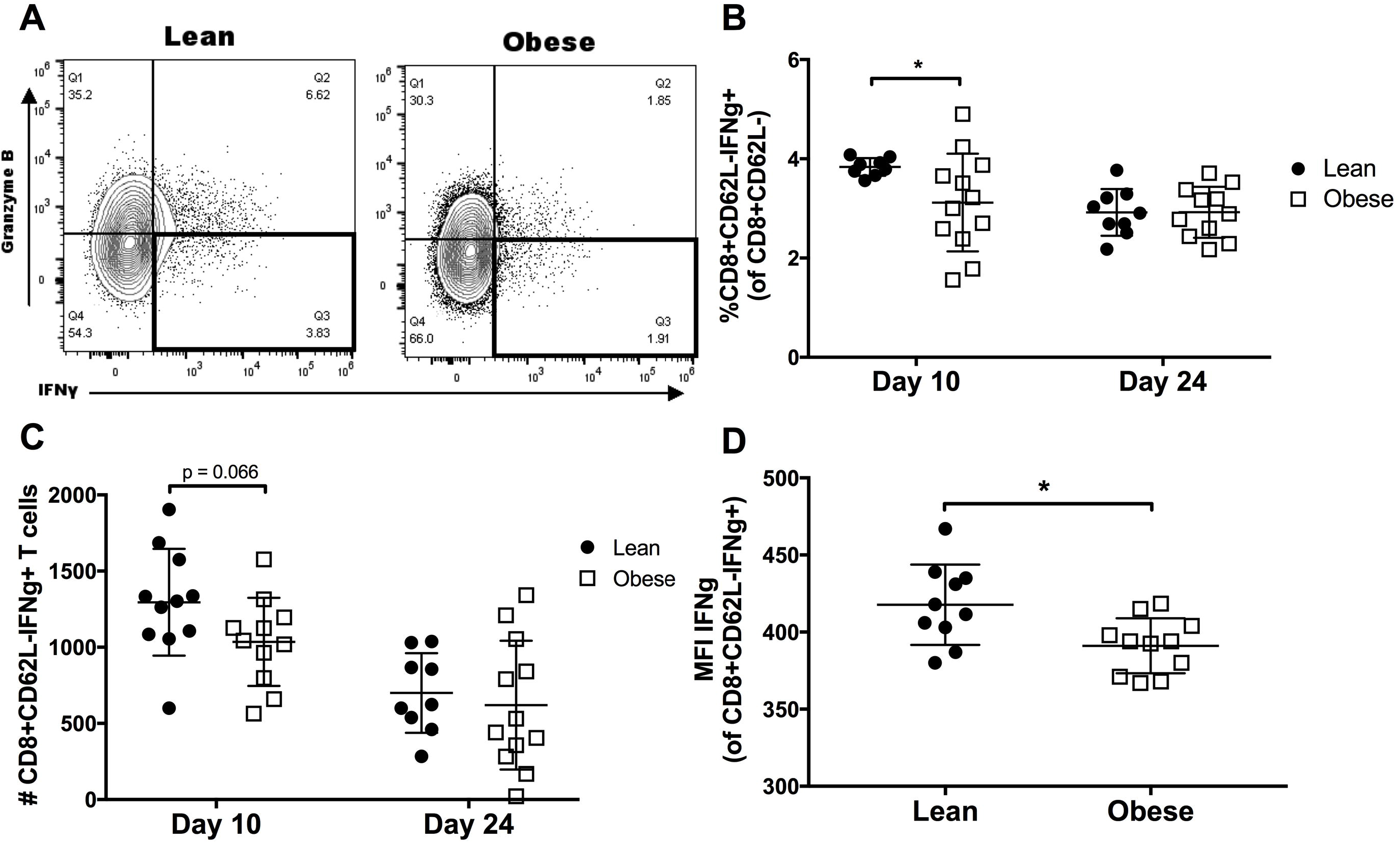
Obesity impairs CD8^+^ T cell production of IFN-γ in lungs of influenza-infected mice. Lungs were harvested, digested, and homogenized into single cell suspension and 1×10^6^ cells were stained for intracellular cytokines in effector T cells by flow cytometry. (A) Representative scatter plot of interferon gamma (IFN-γ) single positive CD8^+^CD62L^−^ T cells at day 10-post infection from lean and obese mice. (B) Frequency and (C) cell number of CD8^+^CD62L^−^GrB^−^IFN ^+^ T cells from day-10 and day-24 post influenza infection in lean (n=11) and obese (n=12) mice. (D) Median fluorescent intensity (MFI) of IFN-γ from CD8^+^CD62L^−^GrB^−^IFNγ^+^ T cells at day 10 post influenza infection. Each dot represents data obtained from one mouse with mean ± SD. Wilcoxon rank sum test was used to compare groups. *p<0.05.

### Obesity decreases naïve and central memory CD8^+^ T cells following influenza clearance

Finally, we assessed if pulmonary T cell metabolic impairments in obesity affects the generation of memory CD8^+^ T cell populations within the lungs of lean and obese mice. At day 10-post primary infection, there were no differences between lean and obese mice in the number (Figure S2A) or percent (Figure S2B) of total, naïve, central memory, or effector memory CD8^+^ T cells. At day 24-post influenza infection, there was a significant decrease in the number of naïve and central memory pulmonary CD8^+^ T cells, with no difference in total, effector memory, or residential memory CD8^+^ T cells (Figure S2C). Finally, lean and obese mice had equivalent percent of total, naïve, central memory, effector memory, and residential memory CD8^+^ T cell populations following influenza clearance at day 24 (Figure S2D).

## DISCUSSION

Influenza remains a serious public health threat, causing approximately 3 to 5 million cases of severe illness and up to 500 000 deaths each year globally (31). Within the United States, seasonal influenza-related illness accounts for 9.2 to 35.6 million cases and anywhere from 140 000 to 710 000 hospitalizations (32). Following the 2009 H1N1 swine flu pandemic, obesity was recognized as an independent risk factor for increased influenza-related morbidity and mortality (33). Since identification as a high-risk population, several studies have described how obesity affects immunity to influenza virus (10, 13, 18, 34–36), however the mechanism(s) responsible remain largely unknown.

Obesity is a complex multisystem metabolic state. Prolonged excess caloric consumption results in greater adipose tissue accumulation, increased presence of circulating pro-inflammatory cytokines, and often insulin or leptin resistance (37). Individual cellular affects of obesity likely contribute to the global pathogenesis of chronic and infectious diseases. DIO animal models mimic the metabolic and morphological features of human obesity (38), thus serving as an appropriate experimental model to dissect the affects of obesity on the immune response to influenza infection.

Foundational work demonstrating the connection between T cell metabolism and function establishes the critical link between cellular glycolysis, glutaminolysis, and fatty acid oxidation to support T cell proliferation, differentiation, function, and survival (39). T cell metabolism is a dynamic process, with the metabolic needs of the cell varying in response to changes in the environment and to activation and stress signals. Activated T cells have a very high metabolic demand, which fuels differentiation, proliferation, and function (24, 40). To meet this increased energy demand, dramatic shifts in the metabolic profile of T cells occur upon activation.

Activated cytotoxic effector CD8^+^ T cells upregulate glycolysis, supporting growth and effector function by forgoing ATP production for nucleotide, amino acids and substrate synthesis used in daughter cell and cytokine production (22, 41, 42). Importantly, glycolytic metabolism is essential for cytokine production (43–45). GAPDH and LDHA have been shown to regulate IFN-γ production through promoter specific direct and independent mechanisms, respectively (28, 46). Further, HIF-1α binding of the IFN-γ promoter has been suggested as a mechanistic link between increased glycolytic metabolism and IFN-γ production by effector T cells (47).

In contrast, naïve and memory CD8^+^ T cells utilize fatty acid oxidation to fuel immune surveillance and survival (48). Manipulation of fatty acid uptake and oxidation alters memory T cell formation and survival (48, 49). Further, reduction of mitochondrial respiratory capacity reduces memory CD8^+^ T cell formation (50). However, much of this foundational work has occurred *in vitro*, emphasizing the need to understand: 1) how a pathogenic challenge *in vivo* influences the metabolic and functional response of CD8^+^ T cells, and importantly, 2) how a metabolic condition like obesity might influence the cellular metabolism and function of pathogen-specific CD8^+^ T cell populations.

Here, we demonstrate for the first time, suppressed oxidative and glycolytic metabolism of effector CD8^+^ T cells in the lungs of influenza infected obese mice. Importantly, pulmonary CD8^+^ T cells exhibited an inability to utilize glucose metabolism when forced using the ATP synthase inhibitor, oligomycin, suggesting obesity impairs CD8^+^ T cell metabolic plasticity.

Additionally, suppressed spare respiratory capacity and ATP production demonstrates an inability of day-10 effector CD8^+^ T cells from obese mice to metabolically respond during influenza infection. Increased uptake of long-chain fatty acid by day-10 effector CD8^+^ T cells suggests metabolic reprogramming in obesity influences nutrient preference during influenza viral infection.

Further, compared to lean mice, DIO mice had reduced effector CD8^+^ T cells making IFN-γ and GrB. Significant reduction in IFN-γ expression by pulmonary effector CD8^+^ T cells, concomitant with impaired CD8^+^ T cell metabolic reprogramming, suggests obesity likely alters the canonical and metabolic signaling programs required for optimal influenza viral immunity. At day 24-post influenza infection, obese mice had decreased numbers of naïve and central memory CD8^+^ T cell populations in the lungs.

Together, these findings establish how diet-induced obesity increases influenza virus pathogenesis through T cell mediated metabolic reprogramming, favoring fatty acid uptake over glycolytic metabolism. This obesity-induced metabolic reprogramming of CD8^+^ T cells results in suppressed T cell function and memory cell generation. As obesity rates continue to rise, and obesity is a recognized independent risk factor for severe influenza outcomes, these results have significant public health implications. Further understanding is required to improve vaccine and therapeutic strategies to reduce the burden of influenza in obese populations.

## LIST OF ABBREVIATIONS

A/PR/8/34: Influenza A Puerto Rico 8 34 H1N1 Virus
ATP: Adenosine Triphosphate
BAL: Bronchoalveolar
BODIPY: Boron-dipyrromethene (4,4-difluoro-4-bora-3a,4a-diaza-s-indacene)
CD: Cluster of Differentiation
CM: Central Memory
DIO: Diet Induced Obese
ECAR: Extracellular Acidification Rate
EDTA: Ethylenediaminetetraacetic Acid
EM: Effector Memory
FCCP: Carbonyl Cyanide 4-(trifluoromethoxy) Phenylhydrazone
GAPDH: Glyceraldehyde 3-phosphate Dehydrogenase
GrB: Granzyme B
H1N1: Hemagglutinin 1 Neuraminidase 1
HAU: Hemagglutination Units
HFD: High Fat Diet
IACUC: Institutional Animal Care and Use Committee
IFN-γ: Interferon gamma
IQR: Interquartile range
LDHA: Lactate Dehydrogenase A
MFI: Median Fluorescent Intensity
OCR: Oxygen Consumption Rate
PBS: Phosphate Buffered Saline
RM: Residential Memory

## AUTHOR CONTRIUBTIONS

Conceptualization, W.D.G., M.A.B., and S.R.S.; Methodology, W.D.G. and A.E.A.; Investigation, W.D.G., A.E.A., Q.S., M.A.B., and S.R.S.; Formal Analysis, W.D.G. and A.E.A.; Writing – Original Draft, W.D.G.; Writing – Review & Editing, W.D.G., A.E.A., M.A.B., and S.R.S.; Funding Acquisition, W.D.G, A.E.A., M.A.B., and S.R.S.; Resources, M.A.B. and S.R.S.; Mentorship, M.A.B. and S.R.S.

## ACKNOWLEDGEMENTS

The authors thank the UNC Flow Cytometry core, supported in part by P30 CA016086 Cancer Center Core Support Grant to the UNC Lineberger Comprehensive Cancer Center. Research reported in this publication was supported in part by the North Carolina Biotech Center Institutional Support Grant 2017-IDG-1025 and by the National Institutes of Health 1UM2AI30836-01. W.D.G. was supported by NIH National Research Service Award T32-DK007686. A.E.A. is supported by the National Science Foundation Graduate Research Fellowship Program under Grant No. 1650116 to AEA. Any opinions, findings, and conclusions or recommendations expressed in this material are those of the author(s) and do not necessarily reflect the views of the National Science Foundation. This work was supported by the National Institutes of Health R01AT008375 (SRS), P30DK056350 (SRS, MAB) and R01DK106090 (NJM).

