## Supplementary Figures for "Metabolic and functional impairment of CD8^+^ T cells from the lungs of influenza-infected obese mice"

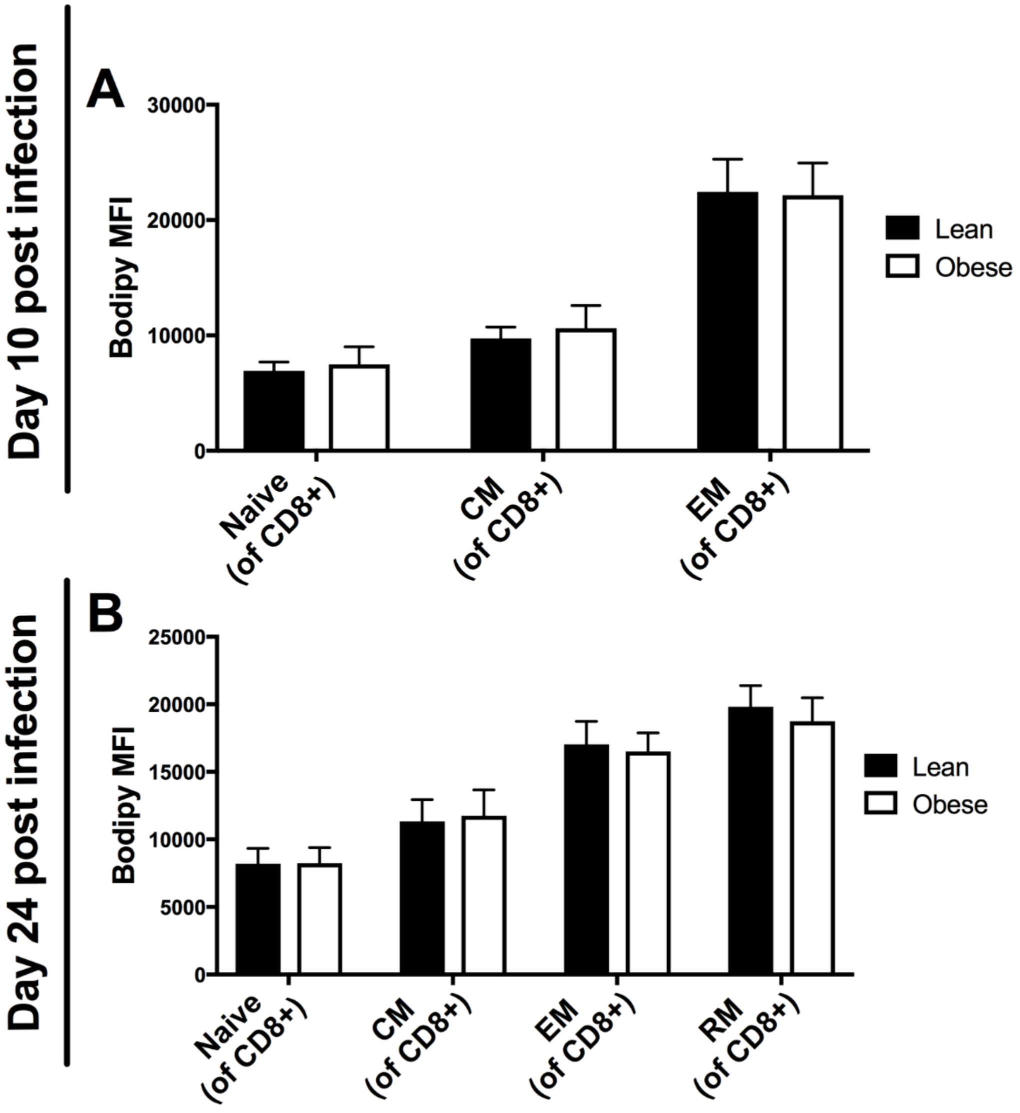


**Figure S1. Related to Figure 4. No difference in fatty acid uptake in naïve, central memory, effector memory, or residential memory $CD8^+$ T cells.**

Lungs were harvested, digested, and homogenized into single cell suspension and $1x10^6$ cells were stained for BODIPY FLC16 uptake in $CD8^+$ populations for naïve ($CD44^-CD62L^+$), central memory ($CD44^+CD62L^+$), effector memory ($CD44^+CD62L^-$), and residential memory ($CD8^+CD69^+CD103^+$) T cells by flow cytometry. (A) BODIPY MFI of naïve, central memory, and effector memory $CD8^+$ T cells at day 10-post influenza infection from lean (n=11) and obese (n=12) mice. (B) BODIPY MFI of naïve, central memory, and effector memory $CD8^+$ T cells at day 24-post influenza infection from lean (n=9) and obese (n=12) mice. Data represent median ± SD. Wilcoxon rank sum test was used to compare groups.

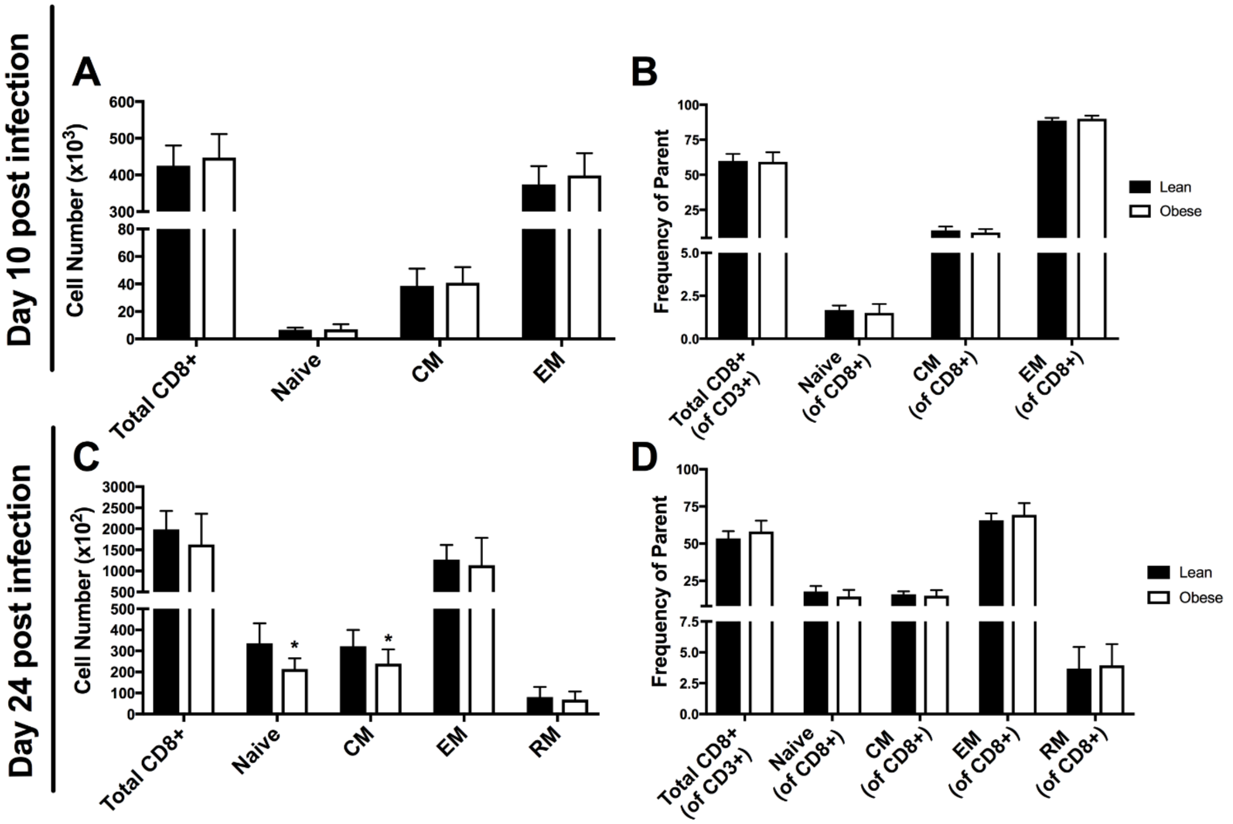


**Figure S2. Obesity impairs number of naïve and central memory CD8$^+$ T cells following influenza clearance.** Lungs were harvested, digested, and homogenized into single cell suspension and 1x10$^6$ cells were stained for total, naïve (CD44$^-$CD62L$^+$), central memory (CD44$^+$CD62L$^+$), effector memory (CD44$^+$CD62L$^-$), and residential memory (CD8$^+$CD69$^+$CD103$^+$) CD8$^+$ T cells by flow cytometry. (A) Number and (B) percent of total, naïve, central memory, and effector memory CD8$^+$ T cells at day 10-post influenza infection from lean (n=12) and obese (n=12) mice. (C) Number and (D) percent of total, naïve, central memory, and effector memory CD8$^+$ T cells at day 24-post influenza infection from lean (n=12) and obese (n=12) mice. Data represent mean ± SD. Wilcoxon rank sum test was used to compare groups. *$p<0.05$.
